# SPIN-CGNN: Improved fixed backbone protein design with contact map-based graph construction and contact graph neural network

**DOI:** 10.1101/2023.07.07.548080

**Authors:** Xing Zhang, Hongmei Yin, Fei Ling, Jian Zhan, Yaoqi Zhou

**Affiliations:** School of Biology and Biological Engineering, South China University of Technology, Guangzhou, China; Institute of Systems and Physical Biology, Shenzhen Bay Laboratory, Shenzhen, China

## Abstract

Recent advances in deep learning have significantly improved the ability to infer protein sequences directly from protein structures for the fix-backbone design. The methods have evolved from the early use of multi-layer perceptrons to convolutional neural networks, transformer, and graph neural networks (GNN). However, the conventional approach of constructing K-nearest-neighbors (KNN) graph for GNN has limited the utilization of edge information, which plays a critical role in network performance. Here we introduced SPIN-CGNN based on protein contact maps for nearest neighbors. Together with auxiliary edge updates and selective kernels, we found that SPIN-CGNN provided a comparable performance in refolding ability by AlphaFold2 to the current state-of-the-art techniques but a significant improvement over them in term of sequence recovery, perplexity, deviation from amino-acid compositions of native sequences, conservation of hydrophobic positions, and low complexity regions, according to the test by unseen structures and “hallucinated” structures. Results suggest that low complexity regions in the sequences designed by deep learning techniques remain to be improved, when compared to the native sequences.

## 1. Introduction

De novo protein design is considered as an inverse problem of de novo protein structure prediction, that is, to find a sequence that would fold into a given structure, instead of predicting its structure for a given sequence. Both problems have been long dominated by energy-based approaches: energy-guided fragment reassembly in the case of protein structure prediction [1] and energy-guided sequence design in the case of protein design [2]. The progress for solving both problems (poor accuracy for de novo structure prediction and low success rate for designed sequences, respectively), however, were hampered by the lack of an accurate energy function to describe the solvent-mediated interactions between amino acid residues of proteins [2, 3].

The energy functions for protein design were typically modified from protein folding studies that can be categorized as molecular mechanics force fields (e.g. EGAD [4]), statistical energy functions (e.g. ABACUS [5, 6]), and mixed statistical, empirical, and physical force fields (e.g. RosettaDesign [7]). More recently, we developed a new protein design technique called OSCAR-design [8] that is based on a purely mathematical scoring function. This scoring function employed series expansion in distance and orientation dependence with mixing coefficients optimized for sequence recovery, sidechain modeling, and loop selection. The optimization effort leads to an average recovery of wild-type sequences at ∼40%, similar to that achieved by the state-of-the-art technique RosettaDesign 3.12 with mixed physical and statistical energy terms, suggesting the bottleneck of an energy-based method.

To avoid an energy function, we developed the first direct prediction of sequences from structures by a simple artificial neural network called SPIN [9] (Sequence Profiles by Integrated Neural networks). By combining local-fragment-derived sequence profiles and nonlocal-energy functions, SPIN achieved a sequence recovery of 30% among 50 test proteins (marked as TS50). SPIN2 [10] employed a deep three-layer neural network and additional structural features to improve SPIN, achieving a higher recovery of 34% on TS50. At the meantime, Qi et al. [11] employed a combination of three neural networks to predict amino-acid type of a center residue from structure fragments constructed with k-nearest neighbors (KNN) residues. With a preset k of 15, this method achieved 34% recovery in the 5-fold cross validation on a dataset constructed on PDB with 30% sequence identity cutoff. These early methods based on MLP (Multi-Layer Perceptrons), which is a basic architecture of artificial neural networks, provided a proof of concept for deep learning-based protein design given a fixed backbone structure.

Rapid advances in deep learning techniques enable the breakthrough of AlphaFold2 [12] and RoseTTAFold [13] in protein structure prediction by avoiding the need for an energy function through the end-to-end learning [14, 15]. During the same period, there is a rapid employment of deep learning in protein design by convolutional neural networks (CNN) such as SPROF [16], DenseCPD [17], and ProDCoNN [18], graph neural networks (GNN) such as GraphTrans [19], GCA [20], GVP [21], AlphaDesign [22], ESM-IF [23], ProteinMPNN [24], PiFold [25], and LM-DESIGN [26], and transformer such as ABACUS-R [27] and ProDesign-LE [28]. These AI-driven backbone-based design substantially improved sequence recovery from 30-34% in MLP based techniques to ∼50-55% by PiFold [25] or LM-DESIGN [26]. Moreover, the methods have developed from fixed backbone design, flexible design based on energy-based structure generation (SCUBA [29]) and sequence-based structure prediction [30, 31] to structure and sequence generators [32–34].

This work focuses on the use of GNN for further improving AI-based fixed-backbone design as it appears to improve over MLP and CNN-based models in the latest development [35]. GraphTrans [19] represented protein backbone structures as a graph, in which residues were represented by node features containing distance and orientation between sequential adjacent *C*_α_atoms and backbone dihedral angles, while inter-residue distances and orientations were encoded as edge features. With an encoder-decoder model constructed with graph attention layers, node features were updated to predict the sequences by sequential autoregressive decoding. GraphTrans achieved 35.82% recovery after training and testing on a dataset set constructed on CATH 4.2 database (marked as CATH4.2). GCA [20] (Global Context Aware generative protein design) improved GraphTrans by appending a global module after the local module (the graph attention layer). It improved the recovery on CATH4.2 to 37.64%. GVP [21] (Geometric Vector Perceptron) decoupled vector and scale information in graph features and proposed a network module to update geometrically sensitive representations. By simply replacing MLP layers employed in GraphTrans with GVP layers, the recovery on CATH4.2 increased to 39.47%. ProteinMPNN [24] improved GraphTrans with three modifications: replacing edge features with interatomic distances between all five atoms (including a virtual *C*_β_atom) on backbones, updating edge features in GNN, and replacing the sequential decoding order with random decoding order in autoregressive decoding. These modifications further improved sequence recovery on CATH4.2 to 45.96%. PiFold [25] introduced virtual atoms determined by backbone position and learnable parameters. Besides, the autoregressive decoding was replaced by one-shot decoding. PiFold achieved 51.66% recovery on CATH4.2 with orders of magnitude efficiency improvement.

The above methods can be further improved by using additional training data, scaling model sizes, and integration with large pretrained models. For example, ESM-IF [23] stack a scaled GVP model with a large transformer model and trained with over 1.2 million of structures predicted by AlphaFold2. The model containing over 142 million parameters significantly improved the recovery of the GVP model in CATH 4.2 dataset from 42.2% to 51.3%. LM-DESIGN [34] used a large protein language model, which was pretrained with over 50 million protein sequences, as a decoder to sampled protein sequences with an encoder from GVP, ProteinMPNN, and PiFold. With about 650 million of additional pretrained parameters, this method brought over 5% improvement to these methods. For the PiFold model reimplemented in this work, we found an increase from 50.22% to 55.65% for the sequence recovery.

These GNN-based methods [19–26], however, utilized k-nearest neighbors (KNN) to construct a graph for feature initialization and local information passing. The KNN graph construction significantly reduces the computational cost from passing information between all node pairs in a graph, since k is much smaller than the length of a sequence in most cases. Moreover, a proper setting of k is expected to prevent the modules for local information passing from overfitting the non-local information. Typically, k is set as 30 because many studies [19, 24] suggested it sufficient for local information passing in GNN-based, fixed-backbone protein design. However, the local structures defined by a fixed number of neighbors might not be capable of handling dense local structures and sparse local structures at the same time.

In this study, we proposed SPIN-CGNN, a deep graph neural network-based method for the fixed backbone design, in which a protein structure graph is constructed with a distance-based contact map. This contact map-based graph construction enables GNN to handle a varied number of neighbors within a preset distance cutoff. In addition, we introduced information of symmetric and second order edges to update edge features. The symmetric edge information enabled information sharing inside an edge pair that connects two nodes. The information on second-order edges is expected to capture high-order interactions between two nodes from their shared neighbors. We found that this SPIN-CGNN achieved 54.81% for sequence recovery. This was achieved by employing a small model of 5.58 million parameters in the absence of pretrained models. Moreover, we further evaluated the method according to amino-acid substitution matrix, sequence complexity, and the deviation of the query structure to the structure predicted by AlphaFold2. These performance measures further support the improvement of SPIN-CGNN over existing state-of-the-art techniques.

## 2. Methods

### 2.1 Dataset

Many GNN-based methods have employed the CATH 4.2 dataset from Ingraham’s [19] to assess their performance. This dataset was constructed by: a) collecting all chains with no more than 500 residues from the CATH 4.2 at 40% sequence identity cutoff; b) randomly split all collected chains into 80%/10%/10% subsets for training, validation, and test; c) removed the entries from these subsets to ensure that there was no overlap in CATH topology (also known as fold) classification between the subsets. Overall, the dataset consisted of a total of 18024, 608, and 1120 structure-sequence pairs in the training, validation, and test subsets from 950, 100, and 150 structural folds, respectively. Thus, there were overlaps in structural folds within each subset.

To avoid performance bias toward specific structural folds within the CATH4.2 test set, we calculated the TM-scores between all test structures and found some structure pairs with high levels of similarity, as illustrated in supplementary Figure S1. Thus, we created a non-structural redundant version of the test set by iteratively removing entries that had the most structural similar entries (TM-score > 0.4) in the test set until no structurally similar entry pairs remained. Ultimately, the resulting CATH4.2-StructNR193 test set contained 193 structure-sequence pairs, after removing 927 out of the original 1120 entries.

To generate another fully independent test set, we performed the following: (a) gathering all PDB structures released after CATH4.2 (April 9^th^, 2019); (b) extracting chains with no less than 30 residues; (c) calculating the TM-score between each chain and all existing entries from the new test set, the CATH4.2 test set, the CATH4.2 validation set, and the CATH4.2 training set; and (d) retaining the chain if the maximum TM-score with all existing entries is no more than 0.4. Finally, we obtained a structural non-redundant test set with 156 entries, which we named as PDB-StructNR156.

Recently, several methods enabled the generation of near-native structures for protein design. We aimed to further evaluate the performance of fixed backbone protein design methods using generated structures. For this purpose, we utilized the ‘Hallucination129’ test set that includes 129 hallucinated structures [31].

### 2.2 Graph Representation

#### Contact Map-based Graph Construction

We defined the neighbors in a graph by contacts between the virtual *C*_β_ atoms with a distance cutoff. The coordinate of virtual *C*_β_ atom of each residue is calculated by

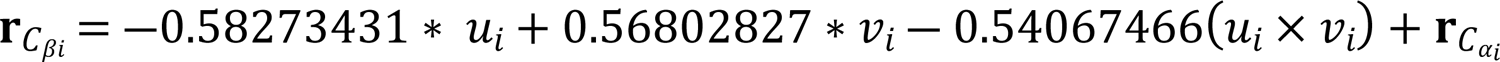

where *u*_i_ = **r**_*C*αi_ ― **r**_Ni_, V_i_ = **r**_*C*i_ ― **r**_*C*αi_. Notably, we removed all self-loops in the contact graph (a residue is not a neighbor of itself). In the contact graph, a residue can receive information from different number of neighbors, depending on the local density around the residue. The edges in a contact graph are symmetric: whenever residue i is a neighbor of residue *j*, residue *j* would be a neighbor of residue i.

#### Node Features

We constructed the node features and edge features as in PiFold [25]. For the node features of residue i, we defined a local coordinate system Q_i_ = [b_i_, n_i_, b_i_ × n_i_] to construct rotation-invariant features, where *u*_i_ is the directional vector from atom *C*_αi_ to atom N_i_(*u*_i_ = **r**_*C*αi_ ― **r**_Ni_), V_i_ is the directional vector from atom *C*_i_ to atom 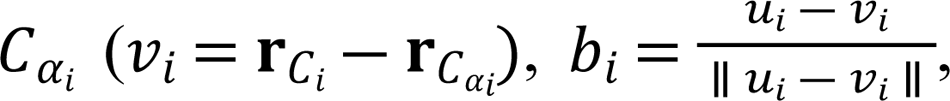 and 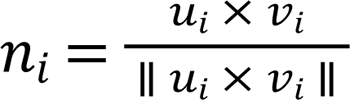. The unit directional vectors from *C*_αi_ to N_i_, *C*_i_, O_i_, and *C*_βi_ in the local coordinate system were collected as node directional features. Bond angles and torsion angles of continuous N_i_, *C*_αi_, and *C*_i_ were encoded as sine and cosine values as node angle features. Specifically, the bond angles included *C*_i―1_-N_i_-*C*_αi_, N_i_-*C*_αi_-*C*_i_, and *C*_αi_-*C*_i_-N_i+1_, where a-b-c denotes the angle between 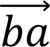 and 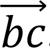. The torsion angles included rotational angles around 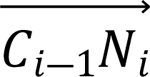, 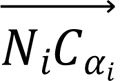, 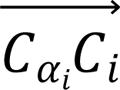 (i.e., ω, ϕ, ψ, respectively). The distances from *C*_αi_ to N_i_, *C*_i_, O_i_, and the virtual *C*_βi_ were encoded with the Gaussian radial basis function (RBF) from ProteinMPNN [24] as node distance features. Finally, the initial node features of residue i were constructed by concatenating all unit vector, angle, and distance features.

#### Edge Features

The edge features of residue *j* to residue i also contain unit vector, angle, and distance features. All interatomic unit directional vectors from five atoms (*C*_α_, *C*, N, O, and the virtual *C*_β_) of residue *j* to those of residue i were calculated and rotated with the local coordinate system Q_i_, in total of 25 edge unit vector features. The interatomic distances between atoms of residue i to atoms of residue *j*, including five main chain atoms (*C*_α_, *C*, N, and O plus the virtual *C*_β_) and three virtual atoms determined by learnable parameters, were collected and encoded with an RBF as 64 edge distance features in total. The coordinate of three virtual atoms {**r**V_i_^1^, **r**V_i_^2^, **r**V_i_^3^} of residue i can be calculated as

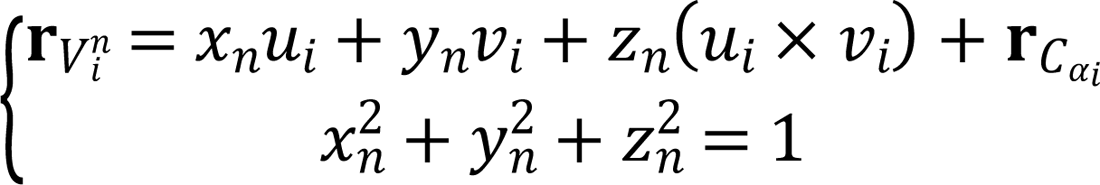

where *u*_i_ = **r**_*C*αi_ ― **r**_Ni_, V_i_ = **r**_*C*i_ ― **r**_*C*αi_, and (*X*_n_, *Y*_n_, *Z*_n_) is a set of learnable parameters of the n^tℎ^virtual atoms. These virtual atoms were employed for capturing complementary information with real atoms. The number of virtual atoms were set as 3 in this study, according to PiFold [25].

The rotation from Q_*j*_ to Q_i_was encoded with the quaternion function as the edge angle features. Additionally, the sequential relative distance (the difference in sequence positions, i ― *j*) from residue *j* to residue i was encoded, with a positional encoding function [36], and append into the edge features. Here, we set the dimension of positional encoding function and RBF as 16. Thus, the total dimensions of node and edge features are 96 and 1119, respectively. Note that any residues with any missing coordinates of atoms *C*_α_, *C*, N, and O will be masked.

### 2.3 Network Architecture

As shown in Figure 1, the graph neural network in SPIN-CGNN was built by stacking 10 Contact Graph Neural Network (CGNN) blocks to fit the message passing in contact map-based graphs. In a CGNN block, edge features were updated first (supplementary Figure S2) and these updated edge features were then utilized to update node features (supplementary Figure S3).

**Figure 1.**
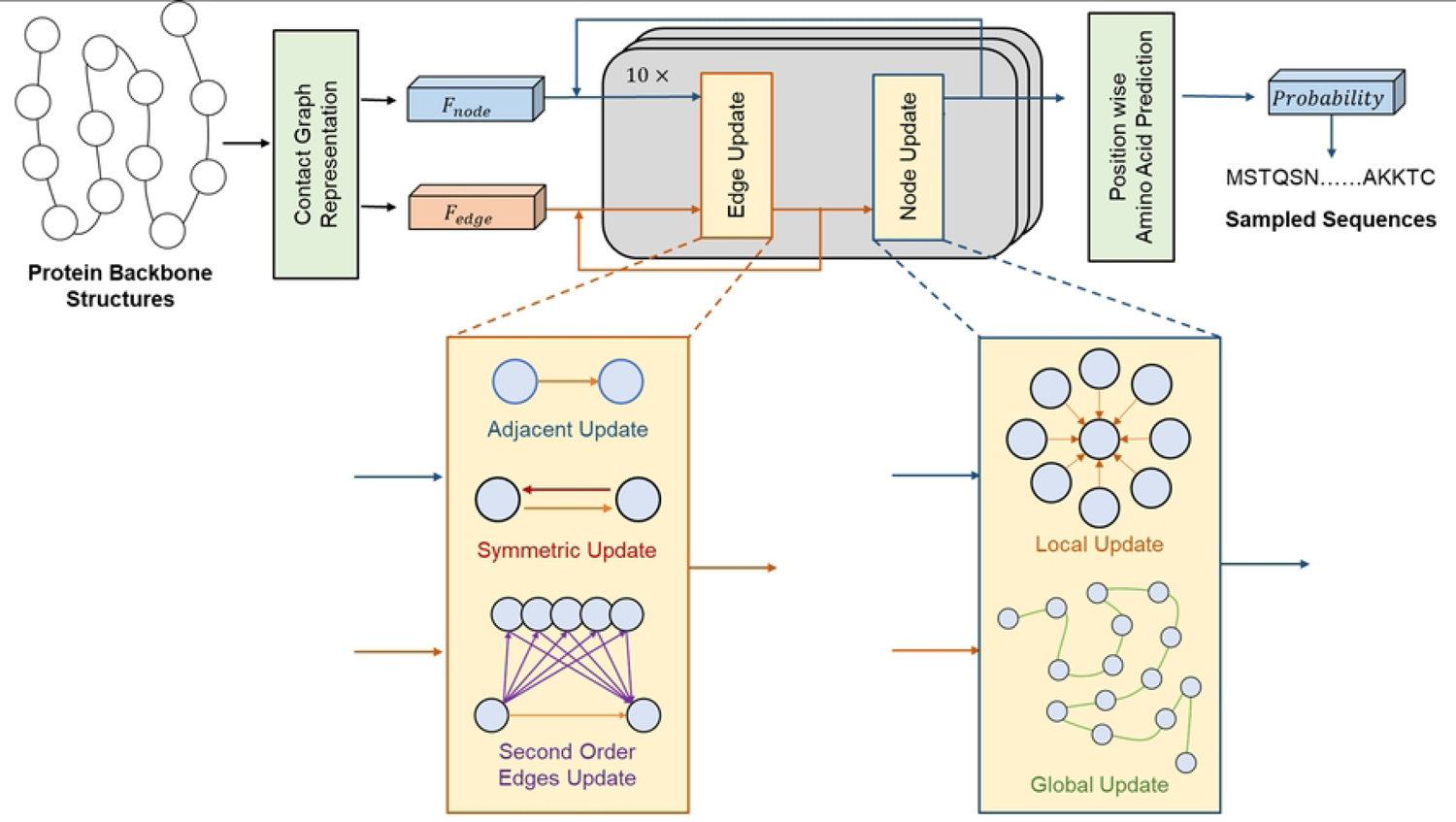
The overall workflow of SPIN-CGNN, which employed a contact-based graph with improved edge and node updates.

#### 2.3.1 Edge update

Updating edge features has been proved to be useful for improving model performance in many related works including ProteinMPNN [24] and PiFold [25]. If we denote the node feature of node *i* in layer *l* as ℎ^l^_i_ and the edge feature from ℎ^l^ to ℎ^l^_i_ to ℎ^l^_j_ as e_ij_, the edge update is performed by simply aggregating the information from adjacent nodes and edge itself:

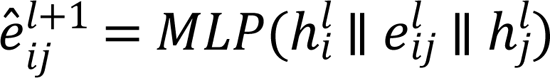

, where e^l+1^ is the edge update by a MLP module, and ∥, represents a concatenate operation.

We further enriched edge updates by edge symmetry. We defined e_i*j*_as the symmetric edge of e_*j*i_in a graph. To ensure that the information passing inside symmetric edge pairs is symmetric, each edge is concatenated with its symmetric edge to produce the symmetric edge information e_symij_^l+1^by:

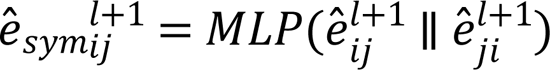

In addition to edge symmetry, we introduced the second-order edges (SOE) by considering the edges associated with their shared neighbors. More specifically, given *N*_i_ representing the neighbor nodes for node i, the shared neighbors of node i and node *j* can be represented as *N*_i_ ∩ *N*_*j*_. For a shared neighbor node n ∈ (*N*_i_ ∩ *N*_*j*_), we defined the combination of e_in_ and e_n*j*_ as the SOE from node i to node *j* though node n. The edge update information from the SOE can be captured with a MLP module from the concatenated feature of ê^l+1^_in_, ê^l+1^_nj_, and ê^l+1^_ij_. By taking the average of all second-order edges of e_i*j*_, we can calculate the SOE information e_soeij_^l+1^ as:

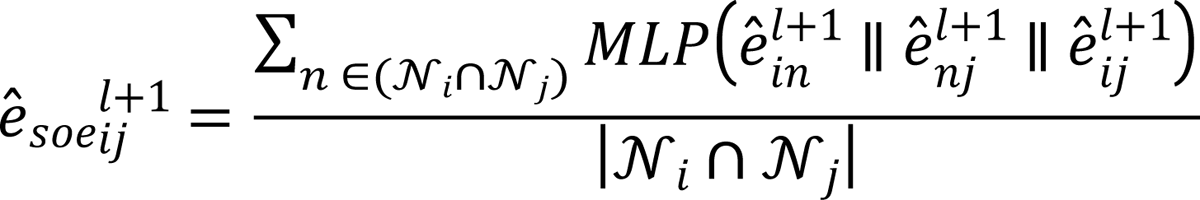

, where |*N*_i_ ∩ *N*_*j*_| is the number of nodes in *N*_i_ ∩ *N*_*j*_. A second MLP module is applied to ê^l+1^_ij_ to specifically extract adjacent information ê_2_^l+1^ from the basic edge update information ê^l+1^_ij_:

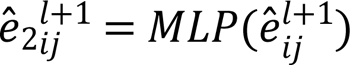

The above updates were merged by a selective kernel module to produce the final edge update (Supplementary Figure S4A), similar to the selective kernel convolution from SK-Net [37]:

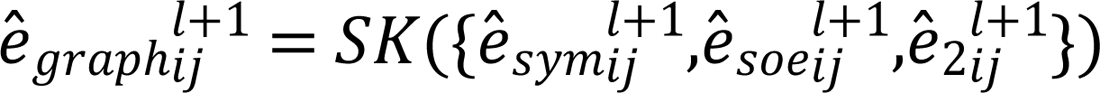

, where *SK* represents selective kernel, and ê^l+1^_graphij_ is the merged edge update information from the graph. To prevent overfitting, the edge information would be updated with dropout, residual connection, and layer normalization:

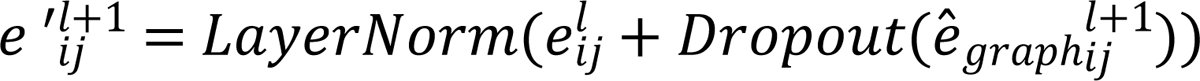

Adding a position-wise feedforward module after attention module has been found to significantly improve the network performance [36]. Therefore, we performed the final update with such typical operation:

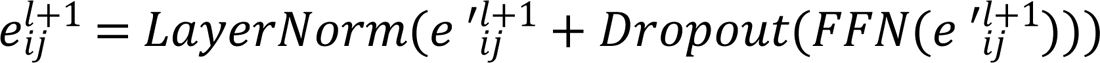

, where *FFN* represent the position-wise feedforward module.

#### 2.3.2 Node update

Node updates in CGNN blocks were performed by integrating local information from neighboring nodes and global information from the whole graph. More specifically, we extract local information by aggregating information from neighboring nodes of the center node with a typical graph-attention module, in which the attention scores were calculated by:

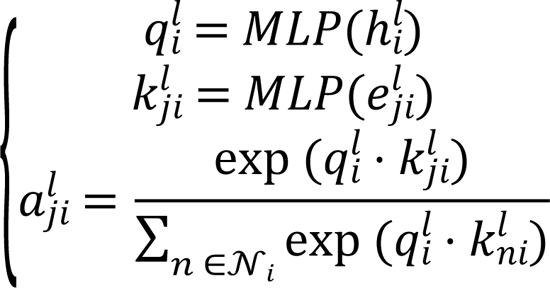

, where *q*^l^_i_ is the query features of node *i, k^l^_ji_* is the key features of e^l^_ji_, and a^l^_ji_ is the attention score of e^l^_ji_ for the local information of node *i*.

We further calculated the value features V^l^_ji_ which were concatenated from edge feature e^l^_ji_, and its adjacent node features ℎ^l^_j_ and ℎ^l^_i_. The summation of neighboring attention-scaled value features yields the local information ℎ_locali_^l+1^

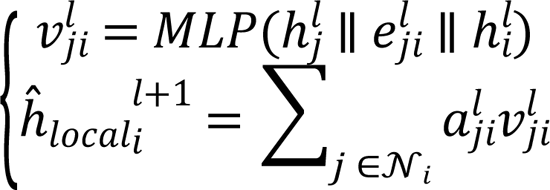

In addition to the local updates, we also used global updates to account for nonlocal interactions. If we define *G* as all nodes in a protein structure graph, we can calculate the global context *G*^l^ by summing scaled value features from all nodes, in which both value feature and attention were calculated from the node feature itself as below:

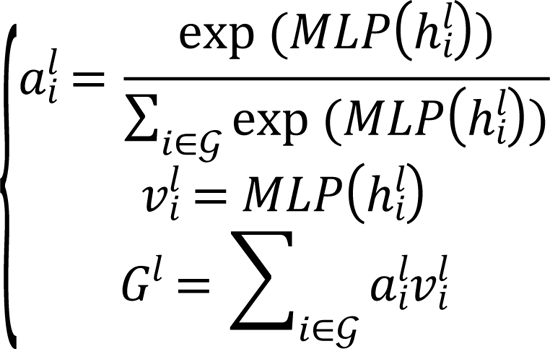

The global update ℎ_globali_^l+1^ was then calculated from the concatenated features of node feature and global context, with an MLP:

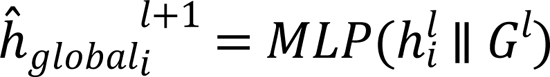

Similarly, a selective kernel module (Figure S4B) was used to merge global with local update information:

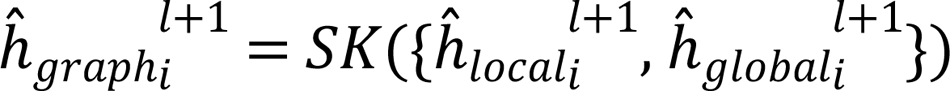

Same as edge update, the node features were updated with the graph update information:

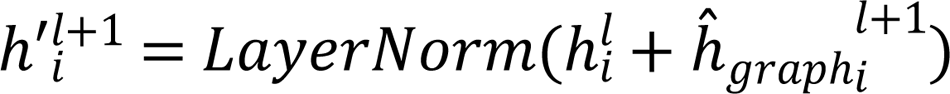

, followed by a *FFN* update:

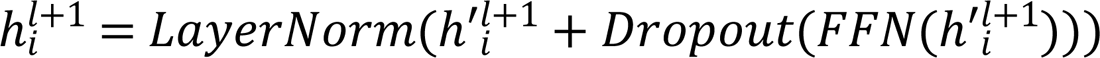

to reduce the possibility of overfitting.

#### 2.3.3 Selective kernel

The selective kernel from SKNet [37] (Selective Kernel Networks) was designed for and has been widely used to merge multi-scale features captured by convolutional kernels with different kernel sizes. In SPIN-CGNN, it was employed to adaptively merge a set of features from different update modules.

Given a feature set ℱ containing n features, a selective kernel simply summarizes all features and squeeze the dimension with an MLP:

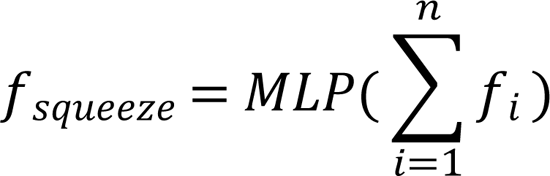

A set of MLPs were employed for the excitation from *f*_squeeze_ to the dimension-wise weight of each feature, and normalized by *Softmax* function:

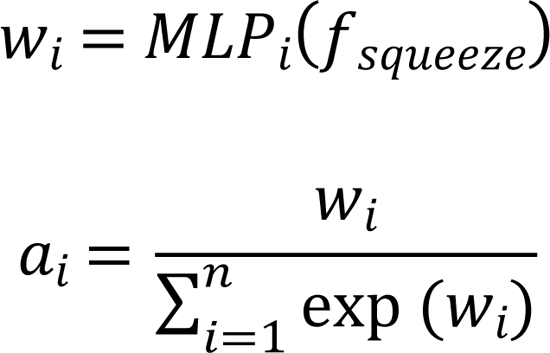

Finally, the merged feature *f*_merge_ is calculated by summarizing all features that are dimension-wisely scaled by the attention:

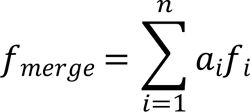

### 2.4 Training

The dimensions of edge and node features were set as 128 for all layers. All models were trained by minimizing the cross entropy between output logits and native sequences for 100 epochs with AdamW optimizer [38]. The learning rate was adjusted according to OneCycle learning rate schedule [39] with a max learning rate of 0.004. We set the drop probability as 0.1 for all dropout operators. Training data were randomly grouped with a maximum batch size of 4096 residues. We employed mixed precision to accelerate the training speed and reduce the GPU memory occupation in all experiments. All other settings followed the default setting of PyTorch.1.13 [40].

### 2.5 Performance Measure

The most widely used criteria for evaluating the methods for fixed backbone design are perplexity and recovery. The perplexity on test set *D* is calculated by exponentiated categorical cross-entropy loss per residue:

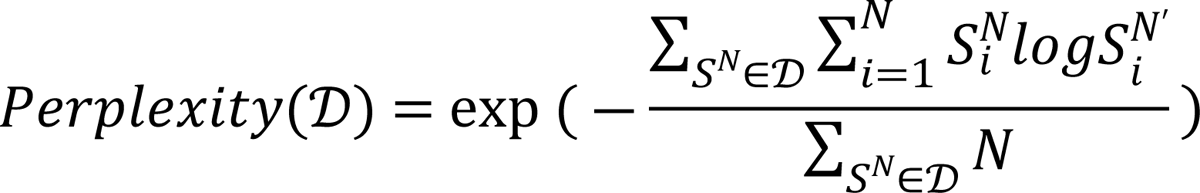

, where *S*^N^ is a sequence with N residues from the test set *D*. *S*^N^_i_ is the *i*-th native residue and *S*^N′^_i_ is the corresponding predicted probability from the model. Perplexity is a measure that accounts for the certainty around the native amino acid residues. Lower perplexity values indicate smaller deviation from native residue types.

Recovery, measuring the ability of the model to reconstruct the native sequence of a protein, is calculated by the percentage of the identity of designed sequences to native sequences:

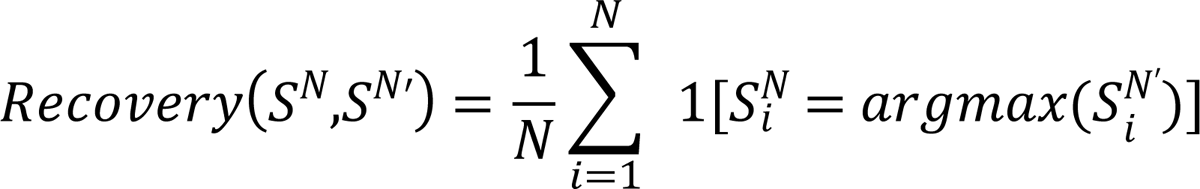

This measure, however, only compared the top ranked prediction and does not reflect fluctuation around the top ranked prediction.

We further examined the frequencies of each amino-acid-residue type given by native sequences and designed sequences. The similarity between native frequencies and predicted frequencies can be measured by the relative deviations:

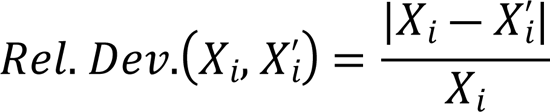

, where *X* is native frequencies and *X*^′^ is the predicted frequencies.

It is known that surface residues are more difficult to recover, we calculated the fraction of surface residues for each target protein. The residue-wise SASA (Solvent Accessible Surface Area) was obtained by BioPython [41]. The SASA for each residue is divided by the maximum allowed solvent accessibility (MaxASA) [42] of the residue type to yield the relative accessible surface area (RSA). Finally, we classified the residues with RSA smaller than 0.2 as core residues, and the others as surface residues as in OSCAR-design [8].

However, the above criteria are based on native sequences. Many sequences can fold into the same structure. Some sequences can fail to fold into a target structure despite high sequence identity to the native sequence because a few mutations may well destabilize the structure. Thus, we also examined low complexity regions, which normally lead to intrinsically disordered regions and the inability to fold into the target structure, with NCBI C++ toolkit [43]. Additionally, we measured the frequencies of amino acid substitutions in the designed sequences from the native sequences by calculation the BLOSUM score and the Pearson correlation coefficient of the confusion matrix with BLOSUM62[44] as the reference of native amino acids substitution. The BLOSUM score is calculated as the summation of BLOSUM62 values of the native amino acid, weighted by the predicted probability. The calculation of confusion matrix followed ESM-IF [23], in which the substitution scores between native sequences and designed sequences were calculated by using the same log odds ratio formula as in the BLOSUM62 substitution matrix. For two amino acid types *X* and *Y*, the substitution score is:

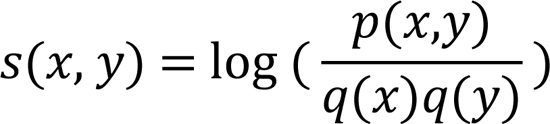

, where *p*(*X*, *Y*) is the jointly likelihood that native amino acid *X* is substituted by predicted amino acid *Y*, *q*(*X*) is the frequencies of amino acid *X* in the native distribution, and *q*(*Y*) is the frequencies of amino acid *Y* in the predicted distribution.

Finally, we performed a test to evaluated the performance of methods by measuring the structure deviations between the target structures and the AlphaFold2-predicted structures for designed sequences. Three criteria were employed including TM-score, RMSD (Root-Mean-Square Deviation), GDT-TS (Global Distance Test – Total Score) on the superposition *C*_α_coordinate. Specifically, the distance cutoff used in GDT-TS is 1, 2, 4, and 8 Å, same as that in CASP.

### 2.6 Method comparison

We employed three methods for comparison including an energy-based method OSCAR-design, and two GNN-based method ProteinMPNN and PiFold. For the energy-based method OSCAR-design, we designed sequences with the default setting.

For deep learning-based methods ProteinMPNN and PiFold, we reimplemented them with the source code and the training setting from their paper. Specifically, we reimplemented ProteinMPNN model with its source code and training setting: negative-log likelihood loss, transformer learning rate schedule, batch size 6000 residues, training epochs 100, Adam optimizer, 30 residue neighbors, no coordinate noise. The median recovery of the reimplemented model tested on CATH4.2 test set is 46.15%, which is consistent to the reported recovery of ProteinMPNN reimplemented model (45.96%) [25]. And we reimplemented PiFold with its source code and training setting: negative-log likelihood loss, OneCycle learning rate schedule, a batch size of 4096 residues, training epochs of 100, Adam optimizer, 30 residue neighbors, and 3 virtual atoms. The median recovery of the reimplemented model tested on CATH4.2 test set is 51.55%, which is also consistent to the reported recovery (51.66%) [25].

## 3. Results

### 3.1 Impact of graph constructions

We examined the impact of using the contact maps at different cutoff distances and compared them against the K-nearest-neighbor graph (k=30, KNN-30) employing the CATH4.2-StructNR193 and the PDB-StructNR156 test sets. Performance was evaluated using perplexity and median recovery. To make a fair comparison, Table 1 compares KNN-30 to CGNN all at without employing CGNN edge information (named as Model 1). The results indicated that KNN-30 has a better performance than CGraph-8 (8Å distance cutoff) for both test sets. However, increasing distance cutoff (from 8Å to 10Å, and then 12 Å) improves over KNN-30 in perplexity and sequence recovery. At 12 Å cutoff, there is ∼1% increase in median sequence recovery, and 2-3% reduction of perplexity from KNN-30 to CGraph-12. We note that even at 12Å cutoff, the average number of neighbors (25 or 29) is still smaller than 30 employed in KNN-30. We fixed the cutoff at 12Å for all subsequent analysis because further increasing the cutoff will only lead to minor improvement at the expense of higher computational requirement.

**Table 1.**
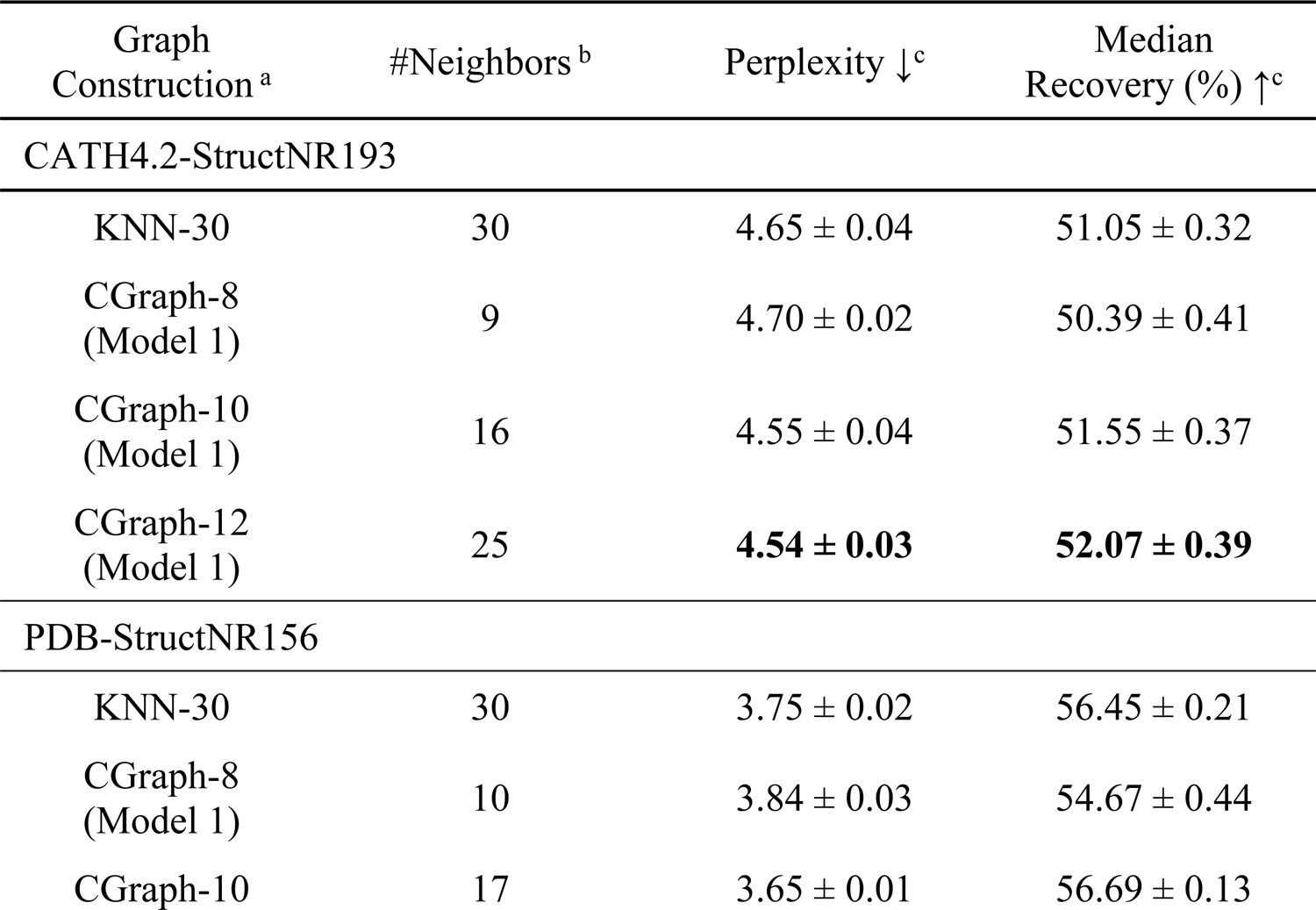
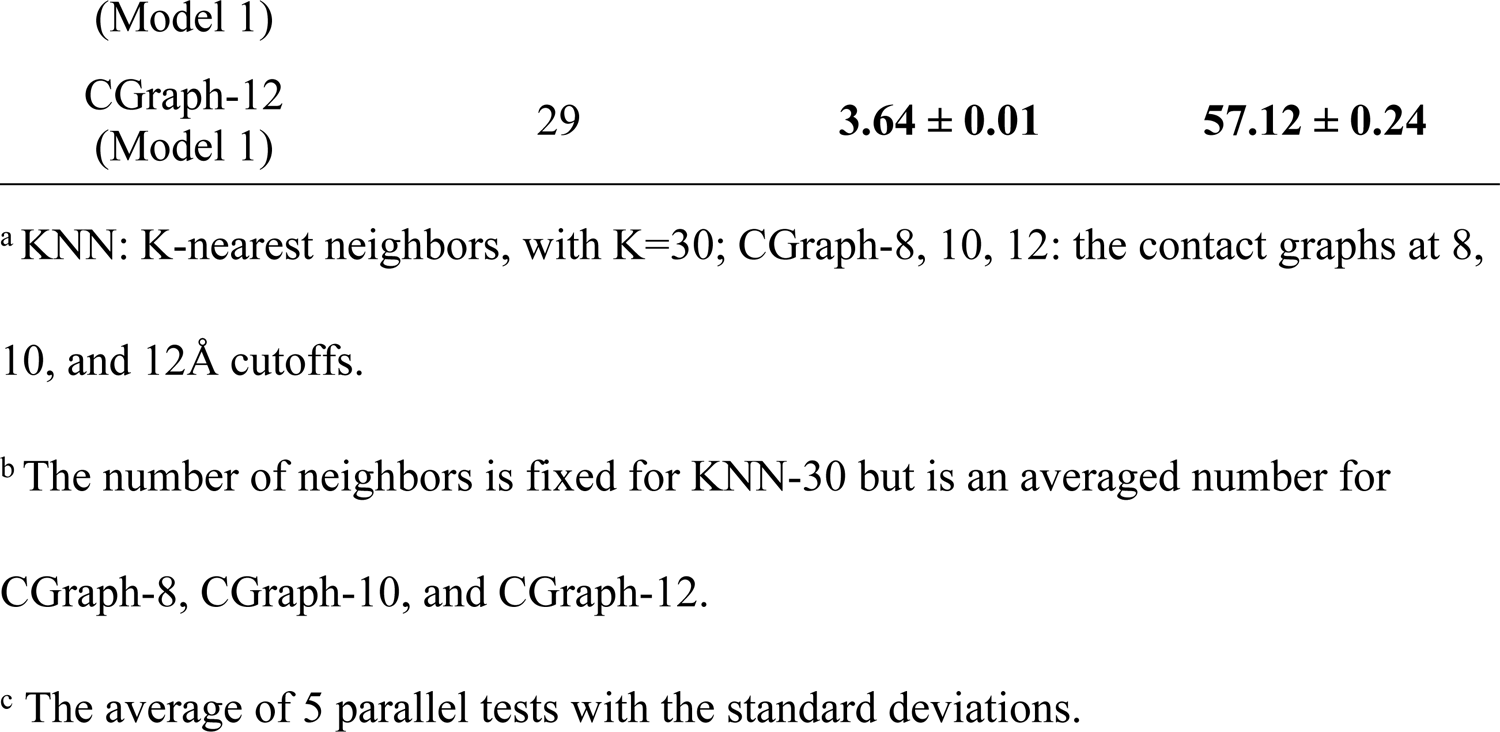
Contact-based versus K-nearest neighbors in the absence of edge information for all methods (Model 1 for SPIN-CGNN) according to perplexity and median sequence recovery for two test datasets (CATH4.2-StructNR193 and PDB-StructNR156).

### 3.2 Ablation test for CGNN edge updates

Table 2 examines the effect of second-order edge (SOE) and symmetric edge updates by constructing Model 2 and Model 3, respectively, as well as the SPIN-CGNN with both SOE and symmetric edge updates. Table 2 shows that without both edge updates Model 1 leads to the worst performance in perplexity and median sequence recovery. Removing SOE also led to statistically significant increase of perplexity and reduction of median recovery from SPIN-CGNN. Although symmetric edge updates do improve the CGNN model when SOE update is absent (from Model 1 to Model 2), it contributes little when the SOE update is performed, indicating that the information captured by symmetric edge updates may have been covered by SOE updates. The overall effect of introducing edge updates is ∼1% increase in sequence recovery and 4-6% reduction in perplexity.

**Table 2.** Impact of CGNN edge updates (symmetric information and second-order edge (SOE) information) according to perplexity and median sequence recovery for two test datasets (CATH4.2-StructNR193 and PDB-StructNR156).

| Model | Sym. | SOE | Perplexity $\downarrow^a$ | Median Recovery (%) $\uparrow^a$ |
| --- | --- | --- | --- | --- |
| CATH4.2-StructNR193 |  |  |  |  |
| Model 1 | × | × | $4.54 \pm 0.03$ | $52.07 \pm 0.39$ |
| Model 2 | √ | × | $4.43 \pm 0.02$ | $52.55 \pm 0.40$ |
| Model 3 | × | √ | $4.36 \pm 0.01$ | **$53.04 \pm 0.31$** |
| SPIN-CGNN | √ | √ | **$4.36 \pm 0.01$** | $52.89 \pm 0.64$ |
| PDB-StructNR156 |  |  |  |  |
| Model 1 | × | × | $3.64 \pm 0.01$ | $57.12 \pm 0.24$ |
| Model 2 | √ | × | $3.53 \pm 0.01$ | $57.50 \pm 0.48$ |
| Model 3 | × | √ | $3.46 \pm 0.02$ | $58.45 \pm 0.50$ |
| SPIN-CGNN | √ | √ | **$3.43 \pm 0.02$** | **$58.54 \pm 0.27$** |

<sup>a</sup> The average of 5 parallel tests with the standard deviations.

### 3.3 Ablation test for selective kernel (SK)

Table 3 examines the effect of the feature integrating module, selective kernels (SKs), compared to average pooling on features to be integrated. Specifically, we constructed three additional models: Model 4, where all SKs in both edge and node updates were replaced by average pooling, Model 5, where only SKs in edge updates were replaced, and Model 6, where SKs in only node updates were replaced. Table 2 indicates that Model 4 had the worst or second-to-the-worst perplexity and median sequence recovery. The cumulative improvement due to the use of SK is 3% reduction in perplexity and 1% increase in sequence recovery.

**Table 3.**
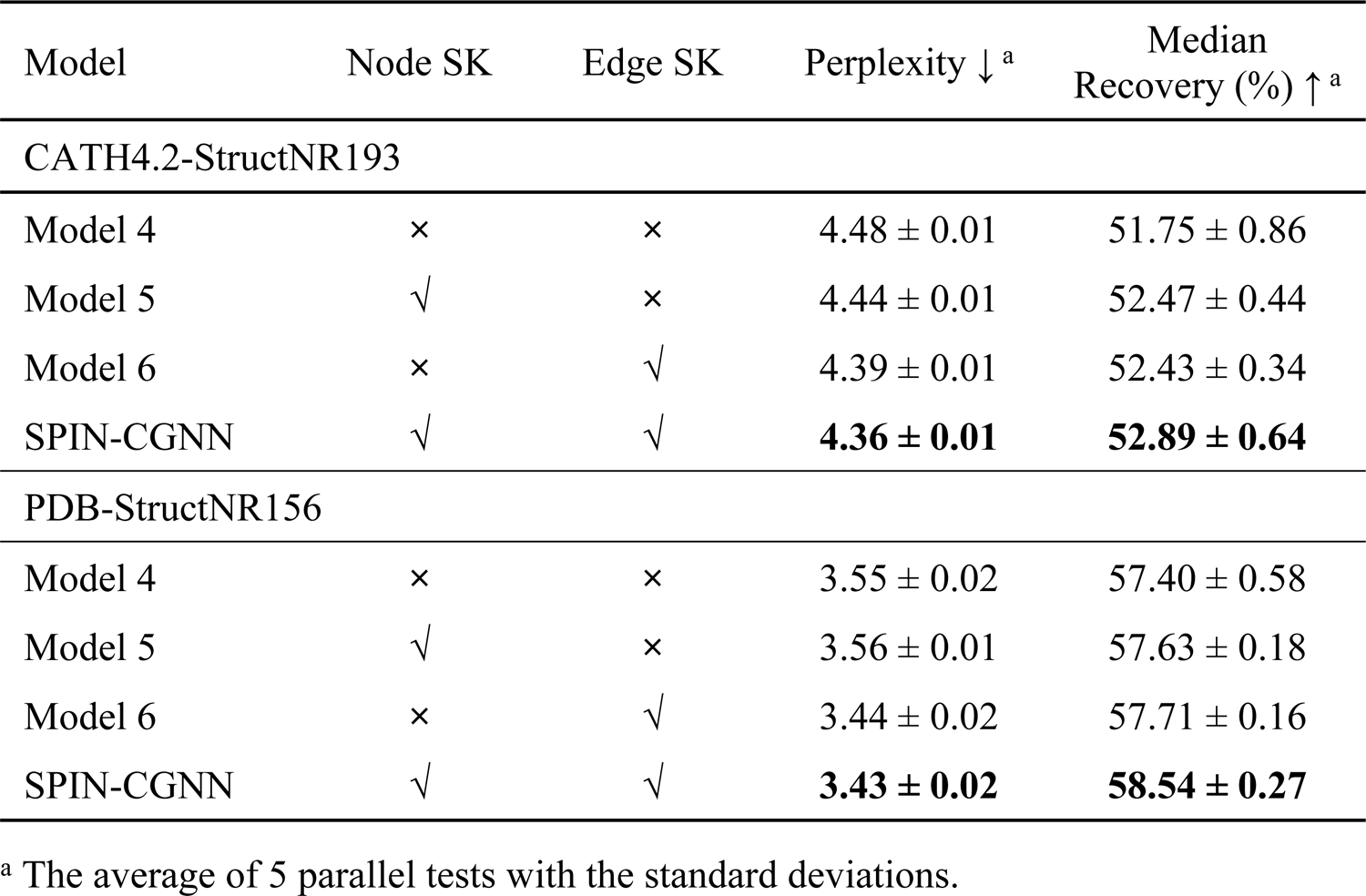
Impact of the use of selective kernels in node update and edge update according to perplexity and median sequence recovery for two test datasets (CATH4.2-StructNR193 and PDB-StructNR156).

We noted that there is a large gap in the performance between CATH4.2-StructNR193 and PDB-StructNR156 test sets. We found that this is mainly because surface residues are less conserved and, thus, harder to recover in computational design and the PDB-StructNR156 test set has 6.5% less surface residues (54.5% versus 61.0%), and thus, with about 5% higher sequence recovery for designed sequences than the CATH4.2-StructNR193 test set. As shown in Supplementary Figure S5, there is an overall similarity in the dependence of recovery on the fraction of surface residues, indicating the robustness of SPIN-CGNN on unseen structures.

### 3.4 Method comparison on the whole CATH4.2 test set

Table 4 compared SPIN-CGNN to a number of other methods for fixed-backbone protein design that employed the same CATH4.2 training, validation and test sets. This is based on the whole test set (rather than structurally non-redundant set) as we do not have the performance for all individual proteins for most methods. As we can see, SPIN-CGNN achieved the best performance in terms of both perplexity and recovery for the whole test set, as well as for two subsets of small and single-chain proteins with 3-4% improvement of recovery and 10-20% improvement in perplexity over the next best PiFold for those methods without a pretrained language model. Compared to LM-DESIGN, which employed the language model for enhancing the method PiFold, our method continues to improve over perplexity by 10% with a slightly lower sequence recovery (1.0%).

**Table 4.**
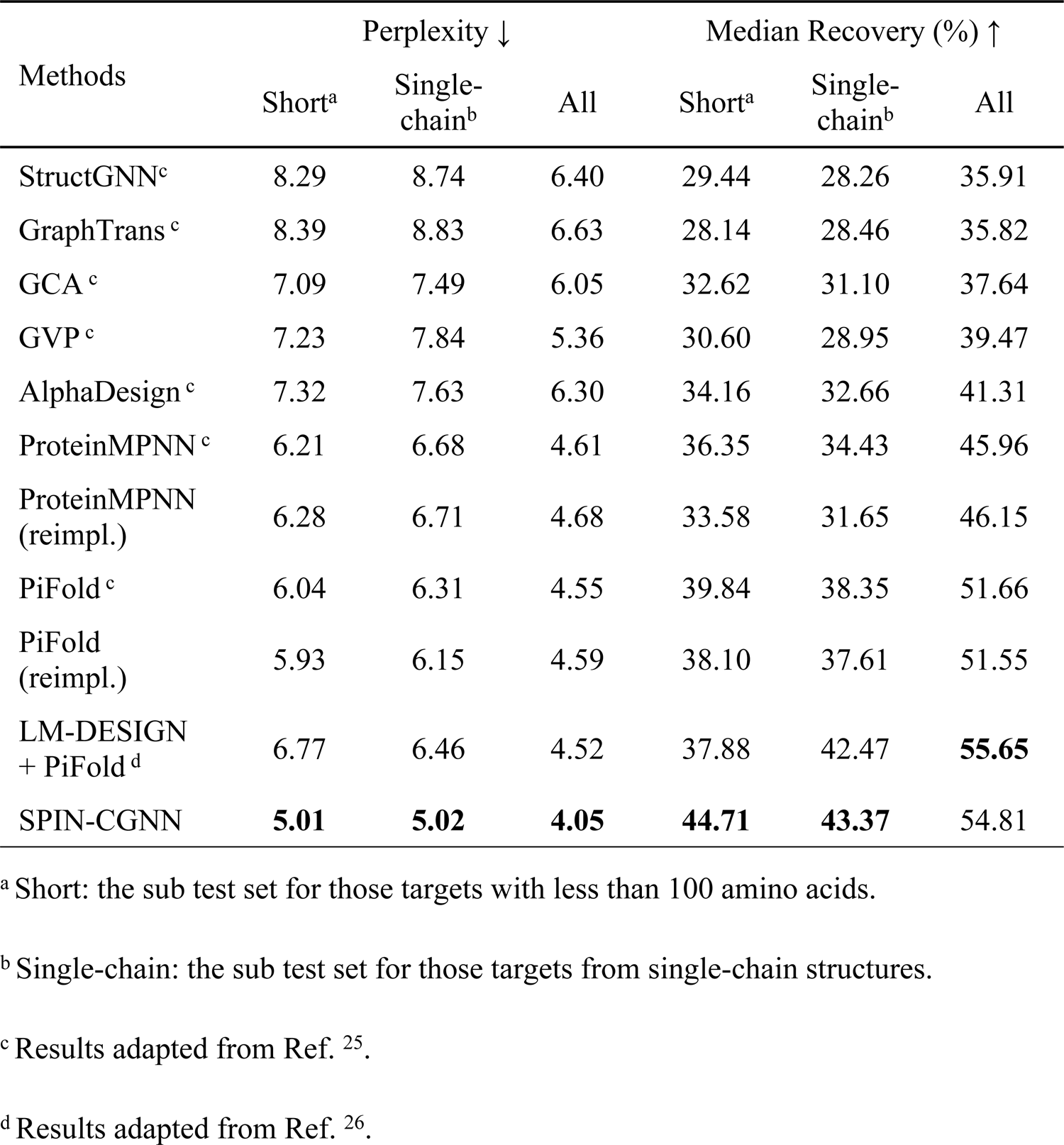
Method comparison on the whole CATH4.2 test set according to perplexity and median native sequence recovery.

### 3.5 Method comparison on structural non-redundant test sets

To confirm that the above improvement by SPIN-CGNN over other methods was not due to biased structural redundancy, Table 5 compared the performance of SPIN-CGNN, OSCAR-design, ProteinMPNN, and PiFold on CATH4.2-StructNR193 and PDB-StructNR156 test sets. Here, we employed OSCAR-design as an example of the energy-based technique, PiFold and ProteinMPNN as examples of modern deep learning models. The results confirmed that SPIN-CGNN has the lowest perplexity (10-15% reduction from the second-best method PiFold, highest sequence recovery (3-4% increase from PiFold).

**Table 5.**
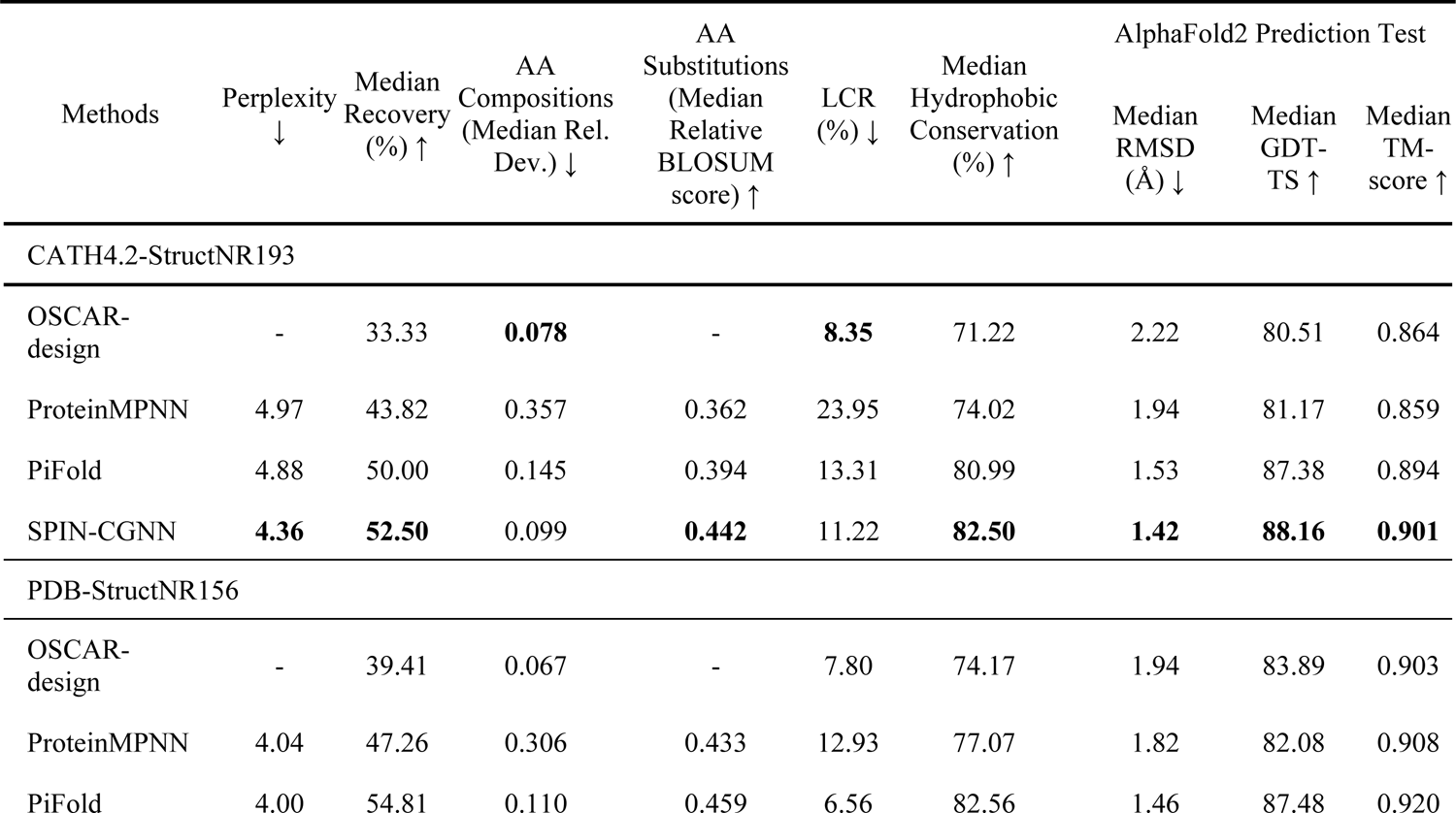
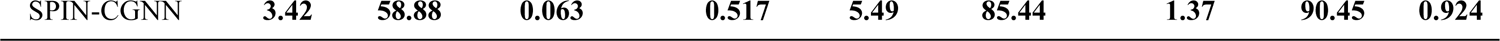
Comparison of sequences designed by SPIN-CGNN, OSCAR-design, ProteinMPNN, and PiFold on CATH4.2-StructNR193 and PDB-StructNR156 test sets according to perplexity, median sequence recovery, median relative deviation of the frequency of amino-acid residue types, the median relative BLOSUM score, fraction of low complexity regions, conservation of hydrophobic and hydrophilic sequence positions, and the difference between refolded and target structures in term of RMSD, GDT-TS and TM-score.

### 3.6 Method comparison on sequence compositions of amino acid residues

Obviously, fluctuation around native sequences (perplexity) and the recovery of native sequences are only one aspect to measure the quality of predicted sequences. The diversity of amino acid residues employed is another measure for designed sequences. A well-designed sequence should take the advantage of the diversity of amino acid residues. Figure 2 compares the frequency of each amino acid residue types employed in native sequences and in the sequences designed by SPIN-CGNN, OSCAR-design, ProteinMPNN, and PiFold, for CATH4.2-StructNR193 (A) and PDB-StructNR156 (B) test sets. There is a large deviation of ProteinMPNN from native frequencies due to its over-employment of A, E, L, and V and under-employment of H, M, Q, R, and W, as shown in Figure 2. The imbalance of residue usages by ProteinMPNN led to the highest median relative deviation of 0.357 (0.306), compared to 0.145 (0.110) by PiFold, 0.099 (0.063) by SPIN-CGNN, and 0.078 (0.067) by OSCAR-design for the CATH4.2-StructNR193 (PDB-StructNR156) dataset (Table 5). Thus, SPIN-CGNN (and OSCAR-design) has much more natural sequence compositions than PiFold and ProteinMPNN (SPIN-CGNN is 32 or 43% better than PiFold, depending on the dataset).

**Figure 2.**
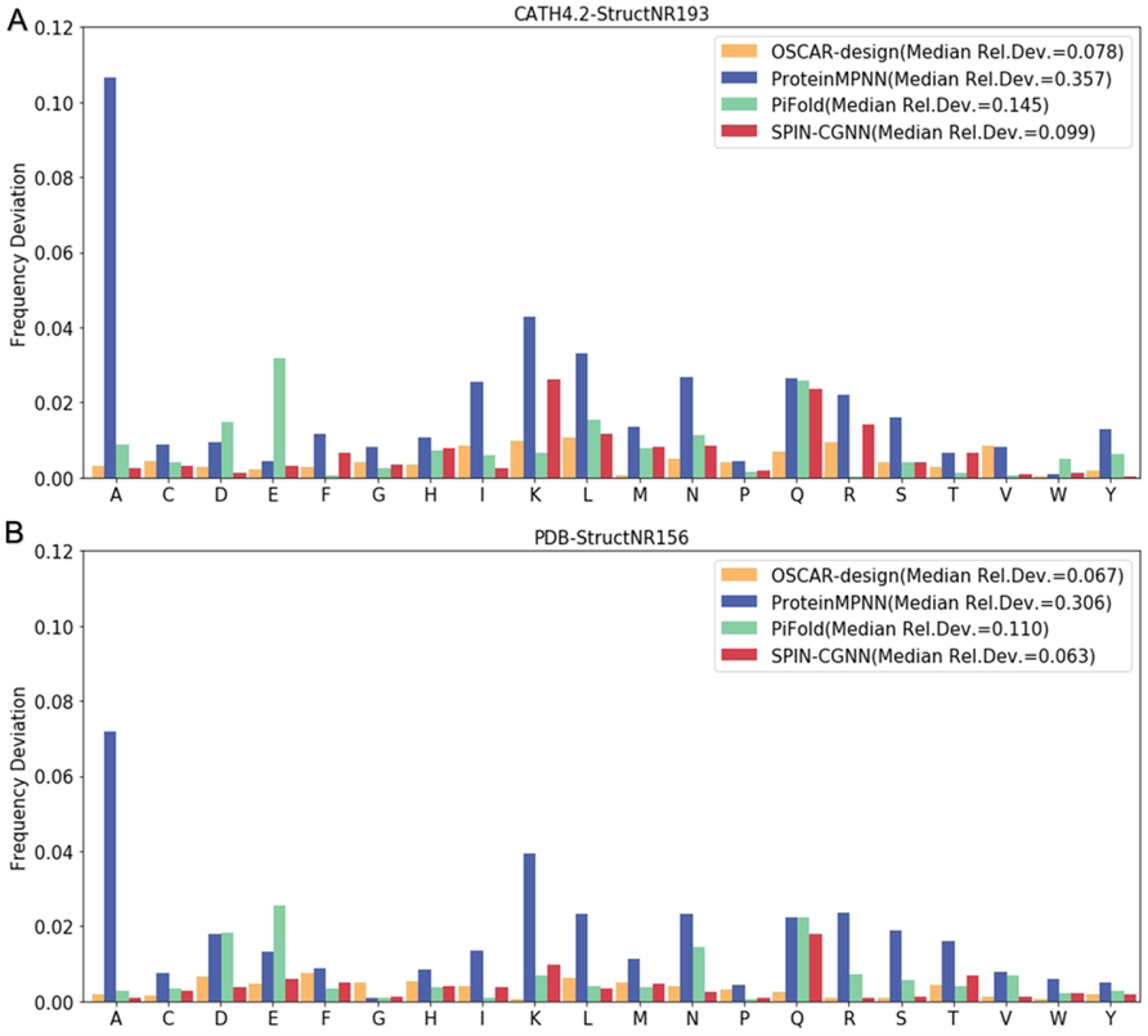
Deviation of the frequency of an amino acid in designed sequences from that in the native sequences by OSCAR-design, ProteinMPNN, PiFold and SPIN-CGNN. (A) CATH4.2-StructNR193 test set and (B) PDB-StructNR156.

### 3.7 Substitutions between amino acids

The phenomenon of amino acid substitutions offers the possibility of different sequences to attain the same target protein structure. This is due to the fact that some positions in the protein structure permit the interchange of amino acids without affecting structural stability. We obtained amino acid substitutions in fixed backbone protein design methods by computing the position-wise confusion matrix between the predicted and native amino acids. Such confusion matrix can be used to compare to BLOSUM62 matrix, that describes the likelihood of amino acid replacements in native sequences. As shown in Figure 3, we can see the confusion matrix of SPIN-CGNN presented a similar pattern to the reference BLOSUM62 matrix: most positive substitutions in BLOSUM62 matrix are also positives values in the confusion matrix of SPIN-CGNN. The Pearson correlation coefficient of SPIN-CGNN on CATH4.2-StructNR193 test set was calculated to be 0.899, compared to 0.884 by OSCAR-Design, (Supplementary Figure S6), 0.839 by ProteinMPNN (Supplementary Figure S7), and 0.890 by PiFold (Supplementary Figure S8). We also obtained the correlation coefficients of these methods on PDB-StructNR156 test set. Similarly, the correlation coefficient given by SPIN-CGNN (0.869) is higher than that of ProteinMPNN (0.853) and PiFold (0.863). Notably, the energy-based method OSCAR-design outperformed all deep learning-based methods with the highest coefficients of 0.888 for this test set.

**Figure 3.**
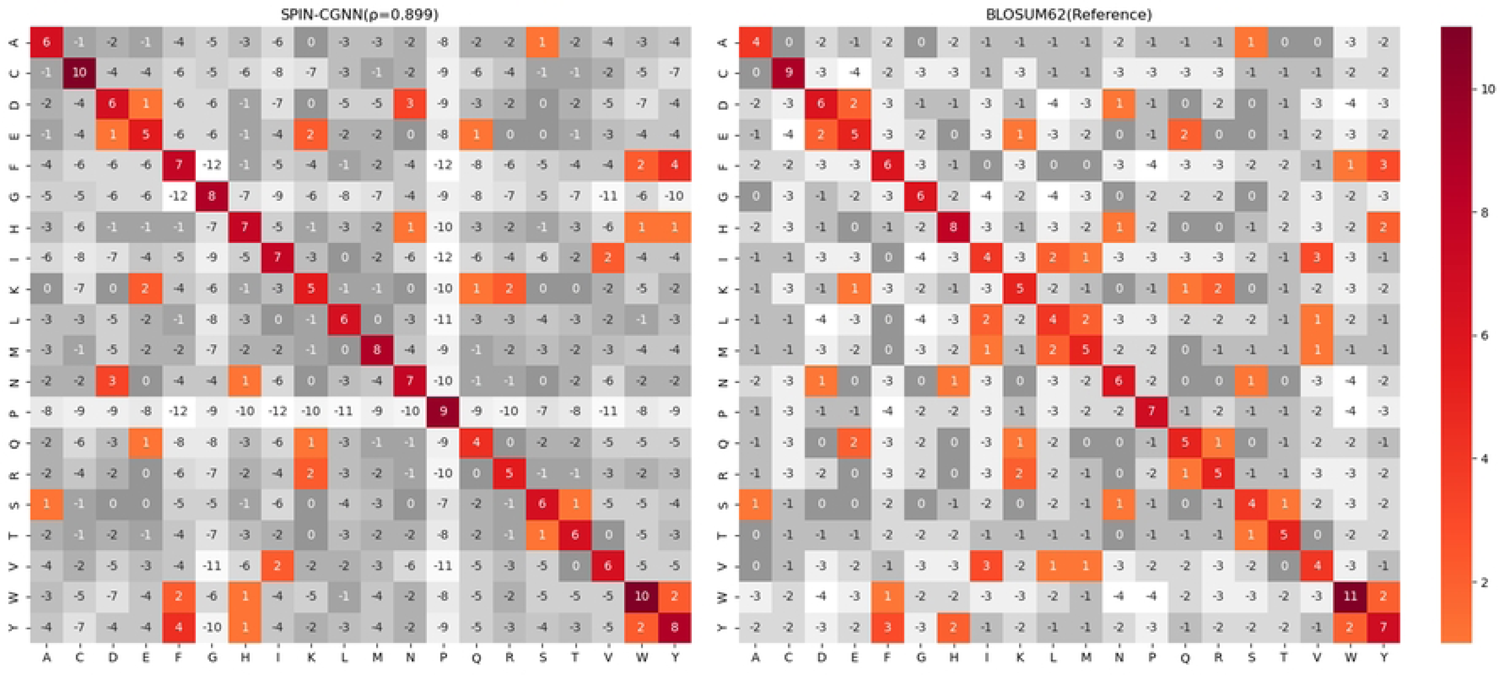
Confusion matrix of SPIN-CGNN in comparison to the reference matrix BLOSUM62 on the CATH4.2-StructNR193 test set. Positive values (colored) indicate substitutions between amino acids. The denoted ρ is the Pearson correlation coefficient between confusion matrix of SPIN-CGNN and BLOSUM62.

We also calculated the BLOSUM score, a summation of BLOSUM62 values weighted by the predicted probability, as a composite metric of perplexity and amino acid substitution. The BLOSUM score of the methods was further normalized by dividing it with the BLOSUM score of the native sequences, where the probabilities of residues were substituted with the one-hot encoded native sequence. As presented in Table 5, SPIN-CGNN outperformed ProteinMPNN and PiFold on both CATH4.2-StructNR193 and PDB-StructNR156 test sets with respect to the median relative BLOSUM score (0.442 / 0.517 for SPIN-CGNN, 0.362 / 0.433 for ProteinMPNN, and 0.394 / 0.459 for PiFold). These results highlight the stronger overall ability of SPIN-CGNN to capture evolution information than other deep learning techniques.

### 3.8 Sequence complexity

The presence and distribution of low-complexity regions (LCR) within protein sequences plays a crucial role in both their structural and functional properties, making it a vital aspect of protein design. Higher LCR fractions in designed sequences as compared to native sequences may result in protein’s structural instability.

The fractions of LCR for native sequences are at 4.12% and 4.27% for the CATH4.2-StructNR193 and the PDB-StructNR156 test sets, respectively. All designed sequences had more LCRs as shown in Table 5. SPIN-CGNN has the lowest (5.5% for the PDB-StructNR156 test set) or the second lowest (11.2% for the CATH4.2-StructNR193, behind OSCAR-design only) fractions of LCRs. Compared to other deep learning techniques, SPIN-CGNN is 1%-2% improvement over PiFold. ProteinMPNN has the worst performance as expected because it over-employed small hydrophobic residues such as A, L, and V (Figure 2).

### 3.9 Hydrophobicity conservation

One important requirement for soluble proteins is that hydrophobic residues should be mostly buried inside the core whereas surface residues are dominated by hydrophilic residues to ensure solubility and prevent hydrophobic aggregation. We examined the conservation of hydrophobic and hydrophilic sequence positions of design sequences by defining hydrophobic (Ile, Leu, Met, Phe, Cys, Trp, Pro, Val, Ala and Gly) and hydrophilic (Ser, Thr, Asn, Gln, Asp, Glu, His, Arg, Lys and Tyr) residue positions according to the native sequence. As Table 5 shows that SPIN-CGNN has the highest conservation in hydrophobicity positions (∼3% over the next best PiFold) for both non-redundant test sets.

### 3.10 Deviation of target structures from the structures predicted by AlphaFold2 based on designed sequences

To further evaluate whether the designed sequences would fold into target structures as expected, we employed AlphaFold2 [12] to predict the structures of designed sequences and measured the root-mean-square deviation (RMSD), global distance test-total score (GDT-TS), and TM-score between predicted structures and target structures. As shown in Table 5, the predicted structures of sequences designed by SPIN-CGNN achieved the smallest median RMSD of 1.42 Å, the greater median GDT-TS of 88.16, and the highest median TM-score of 0.901 on the CATH4.2-StructNR193 test set, comparing to that of OSCAR-design (2.22 Å, 80.51, and 0.864, respectively), ProteinMPNN (1.94 Å, 81.17, and 0.859, respectively), and PiFold (1.53 Å, 87.38, and 0.894, respectively). PiFold and SPIN-CGNN have comparable performance (no statistically significant difference) in term of refoldability by AlphaFold2.

### 3.11 The Hallucination129 test set

The Hallucination129 test set is made of artificially generated structures. As shown in Table 6, OSCAR-design has the lowest fraction of LCR (11%), compared to 15% by SPIN-CGNN, 17% by PiFold, and 33% by ProteinMPNN. For the difference between target structures and AlphaFold2-predicted structures (Figure 4), only ProteinMPNN has highly significant worse performance on GDT-TS and TM-score, and RMSD from PiFold, SPIN-CGNN, and OSCAR-design. The deviations of refolded from native structures given by PiFold, SPIN-CGNN, and OSCAR-design are statistically similar to each other. Adding Gaussian noise of 0.02 Å standard deviation to coordinates did not lead to further improvement of deep learning techniques on artificially generated structures, unlike a previous report [24] (Supplementary Table S1).

**Figure 4.**
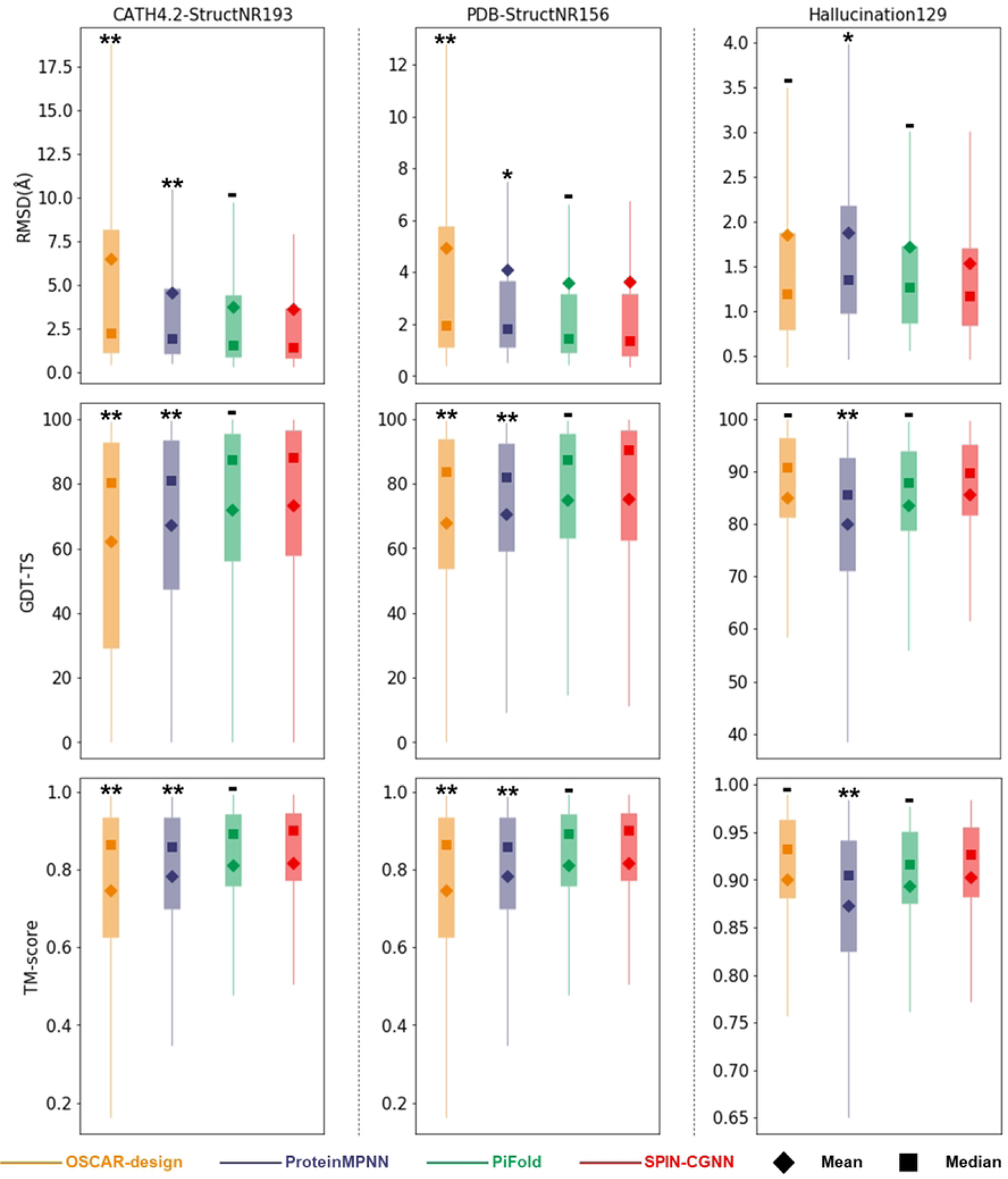
Deviations of the structures of designed sequences predicted by AlphaFold2 from their respective target structures on three separate test sets from left to right panels (CATH4.2-StructNR193, PDB-StructNR156, and Hallucination129 test set) evaluated according to RMSD (Å), GDT-TS, and TM-score (from top to bottom panels). The statistical significance of the difference of a given method to SPIN-CGNN was marked with ‘**’ for highly statistically significant (p-value<0.01), ‘*’ for statistically significant (0.01<p-value<0.05), and ‘-’ for not statistically significant (p-value>0.05). Specific p-values are presented in Supplementary Table S2.

**Table 6.** Comparison of sequences designed by SPIN-CGNN, OSCAR-design, ProDesign-LE, ProteinMPNN, and PiFold on the Hallucination129 test set according to the fraction of Low-Complexity Regions (LCR) and the difference between refolded and target structures in term of RMSD, GDT-TS and TM-score.

| Methods | LCR (%) ↓ | AlphaFold2 Prediction Test |  |  |
| --- | --- | --- | --- | --- |
|  |  | Median RMSD (Å) ↓ | Median GDT-TS ↑ | Median TM-score ↑ |
| ProDesign-LE<sup>28</sup> | - | - | - | 0.83 (Mean) |
| OSCAR-design | **11.0** | 1.19 | **90.75** | **0.932** |
| ProteinMPNN | 32.96 | 1.35 | 85.75 | 0.905 |
| PiFold | 17.19 | 1.27 | 88.00 | 0.917 |
| SPIN-CGNN | 14.91 | **1.17** | 89.75 | 0.927 |

## 4. Discussion

We proposed SPIN-CGNN, a deep learning-based method for fixed backbone protein design. We showed that introducing contact-map, second-order edge updates, and selective kernels cumulatively improved perplexity, sequence recovery, amino-acid composition, amino-acid substitution, low complexity regions, and conservations of hydrophobic/hydrophilic positions, over the most recently developed deep learning-based methods ProteinMPNN and PiFold. Refolding of designed sequences by AlphaFold2 indicates that the structures produced by SPIN-CGNN are comparable to those by PiFold but significantly closer to target structures than ProteinMPNN.

Interestingly, a recently developed energy-based technique, OSCAR-design, produced comparable performance to PiFold and SPIN-CGNN for structures refolded by AlphaFold2 for the Hallucination129 test set. Depending on the datasets, OSCAR-design can have the closest amino-acid composition and low complexity region to the native composition compared to ProteinMPNN, PiFold and SPIN-CGNN, indicating that there is something that deep learning techniques can be improved further.

When we detected the structural redundancy within the CATH4.2 test set, we also found structure pairs with high structure similarity (TM-score>0.4) between all three CATH4.2 subsets (training, validation, and test). To reduce the potential of overfitting, we removed all structures in the training set that have TM-score >0.4 with any structures in the validation and test sets. This led to a much smaller training set of 9311 structures, compared to 18024 proteins in the original set. This new training set, however, led to a poorer performance for those unseen structures. For example, the median TM-score refolded by AlphaFold2 was reduced from 0.924 trained by the whole training set to 0.823 trained by the new training set for SPIN-CGNN for the PDB StructNR156. The similar behavior was observed for PiFold and ProteinMPNN. Thus, we employed the whole training set as it improves the generalizability over the smaller training set.

We would like to emphasize the importance of using different measures to computationally assess the designed sequences. This is because native sequence recovery and the deviation from the native sequence (perplexity) only reflect one aspect of designed sequences. High sequence identity does not assure foldability as a few mutations are often found sufficient to disrupt structural stability. Moreover, too many hydrophobic residues on the protein surface will lead to insoluble and aggregated proteins and low complexity in sequences often leads to intrinsically disordered regions. In addition, because of over-optimization of sequences to the target structure, the refoldability by AlphaFold2 is not that sensitive to the sequences given by different methods (Table 6). In this paper, we showed that the fraction of low complexity regions (11% by SPIN-CGNN, the best in the deep learning methods for CATH4.2-StructNR193) remain much higher than native sequences (4%), indicating the room for further improvement.

It is noted that the number of parameters employed by SPIN-CGNN is 5.58 million, comparable to 4.13 million by PiFold. It has a similar inference time as PiFold. For a 500-residue protein, the inference time is 0.09 second by SPIN-CGNN, compared to 0.03 second by PiFold, 0.83 by ProteinMPNN. Code Availability: source code and datasets are available at https://github.com/EricZhangSCUT/SPIN-CGNN.

## Funding

This work was supported by National Key Research and Development Program of China [NO.2021YFF1200400] and Major Program of Shenzhen Bay Laboratory [S201101001].

## Acknowledgement

We acknowledge Dongbo Bu for his suggestion of the method for the BLOSUM score calculation. The work was done by using the supercomputing facility of the Shenzhen Bay Laboratory.

## Notes

### Competing Interest Statement

The authors have declared no competing interest.

## Reference

1. Bujnicki J M. Protein-structure prediction by recombination of fragments. Chembiochem, 2006, 7(1): 19–27.

2. Li Z, Yang Y, Zhan J, et al. Energy functions in de novo protein design: current challenges and future prospects. Annual review of biophysics, 2013, 42: 315–335.

3. Kuhlman B, Bradley P. Advances in protein structure prediction and design. Nature Reviews Molecular Cell Biology, 2019, 20(11): 681–697.

4. Pokala N, Handel T M. Energy functions for protein design: adjustment with protein–protein complex affinities, models for the unfolded state, and negative design of solubility and specificity. Journal of molecular biology, 2005, 347(1): 203–227.

5. Xiong P, Wang M, Zhou X, et al. Protein design with a comprehensive statistical energy function and boosted by experimental selection for foldability. Nature communications, 2014, 5(1): 5330.

6. Xiong P, Hu X, Huang B, et al. Increasing the efficiency and accuracy of the ABACUS protein sequence design method. Bioinformatics, 2020, 36(1): 136–144.

7. Leaver-Fay A, Tyka M, Lewis S M, et al. ROSETTA3: an object-oriented software suite for the simulation and design of macromolecules. Methods in enzymology. Academic Press, 2011, 487: 545–574.

8. Liang S, Li Z, Zhan J, et al. De novo protein design by an energy function based on series expansion in distance and orientation dependence. Bioinformatics, 2022, 38(1): 86–93.

9. Li Z, Yang Y, Faraggi E, et al. Direct prediction of profiles of sequences compatible with a protein structure by neural networks with fragment-based local and energy-based nonlocal profiles. Proteins: Structure, Function, and Bioinformatics, 2014, 82(10): 2565–2573.

10. O’Connell J, Li Z, Hanson J, et al. SPIN2: Predicting sequence profiles from protein structures using deep neural networks. Proteins: Structure, Function, and Bioinformatics, 2018, 86(6): 629–633.

11. Wang J, Cao H, Zhang J, et al. Computational protein design with deep learning neural networks. Scientific reports, 2018, 8(1): 1–9.

12. Jumper J, Evans R, Pritzel A, et al. Highly accurate protein structure prediction with AlphaFold. Nature, 2021, 596(7873): 583–589.

13. Baek M, DiMaio F, Anishchenko I, et al. Accurate prediction of protein structures and interactions using a three-track neural network. Science, 2021, 373(6557): 871–876.

14. AlQuraishi M. End-to-end differentiable learning of protein structure. Cell systems, 2019, 8(4): 292–301. e3.

15. Ingraham J, Riesselman A, Sander C, et al. Learning protein structure with a differentiable simulator. International Conference on Learning Representations. 2019.

16. Chen S, Sun Z, Lin L, et al. To improve protein sequence profile prediction through image captioning on pairwise residue distance map. Journal of chemical information and modeling, 2019, 60(1): 391–399.

17. Qi Y, Zhang J. DenseCPD: improving the accuracy of neural-network-based computational protein sequence design with DenseNet. Journal of chemical information and modeling, 2020, 60(3): 1245–1252.

18. Zhang Y, Chen Y, Wang C, et al. ProDCoNN: Protein design using a convolutional neural network. Proteins: Structure, Function, and Bioinformatics, 2020, 88(7): 819–829.

19. Ingraham J, Garg V, Barzilay R, et al. Generative models for graph-based protein design. Advances in neural information processing systems, 2019, 32.

20. Tan C, Gao Z, Xia J, et al. Generative de novo protein design with global context. arXiv preprint arXiv:2204.10673, 2022.

21. Jing B, Eismann S, Suriana P, et al. Learning from protein structure with geometric vector perceptrons. arXiv preprint arXiv:2009.01411, 2020.

22. Gao Z, Tan C, Li S Z. Alphadesign: A graph protein design method and benchmark on alphafolddb. arXiv preprint arXiv:2202.01079, 2022.

23. Hsu C, Verkuil R, Liu J, et al. Learning inverse folding from millions of predicted structures. International Conference on Machine Learning. PMLR, 2022: 8946–8970.

24. Dauparas J, Anishchenko I, Bennett N, et al. Robust deep learning–based protein sequence design using ProteinMPNN. Science, 2022, 378(6615): 49–56.

25. Gao Z, Tan C, Li S Z. PiFold: Toward effective and efficient protein inverse folding. arXiv preprint arXiv:2209.12643, 2022.

26. Zheng Z, Deng Y, Xue D, et al. Structure-informed Language Models Are Protein Designers. bioRxiv, 2023: 2023.02. 03.526917.

27. Liu Y, Zhang L, Wang W, et al. Rotamer-free protein sequence design based on deep learning and self-consistency. Nature Computational Science, 2022, 2(7): 451–462.

28. Huang B, Fan T, Wang K, et al. Accurate and efficient protein sequence design through learning concise local environment of residues. Bioinformatics, 2023, 39(3): btad122.

29. Huang B, Xu Y, Hu X, et al. A backbone-centred energy function of neural networks for protein design. Nature, 2022, 602(7897): 523–528.

30. Wang J, Lisanza S, Juergens D, et al. Scaffolding protein functional sites using deep learning. Science, 2022, 377(6604): 387–394.

31 Anishchenko I, Pellock S J, Chidyausiku T M, et al. De novo protein design by deep network hallucination. Nature, 2021, 600(7889): 547–552.

32. Anand N, Achim T. Protein structure and sequence generation with equivariant denoising diffusion probabilistic models. arXiv preprint arXiv:2205.15019, 2022.

33. Watson J L, Juergens D, Bennett N R, et al. Broadly applicable and accurate protein design by integrating structure prediction networks and diffusion generative models. bioRxiv, 2022: 2022.12. 09.519842.

34. Madani A, Krause B, Greene E R, et al. Large language models generate functional protein sequences across diverse families. Nature Biotechnology, 2023: 1–8.

35 陈志航, 季梦麟, 戚逸飞. 人工智能蛋白质结构设计算法研究进展. 合成生物学, 2023: 1.

36. Vaswani A, Shazeer N, Parmar N, et al. Attention is all you need. Advances in neural information processing systems, 2017, 30.

37. Li X, Wang W, Hu X, et al. Selective kernel networks. Proceedings of the IEEE/CVF conference on computer vision and pattern recognition. 2019: 510–519.

38. Loshchilov I, Hutter F. Decoupled weight decay regularization. arXiv preprint arXiv:1711.05101, 2017.

39. Smith L N, Topin N. Super-convergence: Very fast training of neural networks using large learning rates. Artificial intelligence and machine learning for multi-domain operations applications. SPIE, 2019, 11006: 369–386.

40. Paszke A, Gross S, Massa F, et al. Pytorch: An imperative style, high-performance deep learning library. Advances in neural information processing systems, 2019, 32.

41. Cock P J A, Antao T, Chang J T, et al. Biopython: freely available Python tools for computational molecular biology and bioinformatics. Bioinformatics, 2009, 25(11): 1422.

42. Tien M Z, Meyer A G, Sydykova D K, et al. Maximum allowed solvent accessibilites of residues in proteins. PloS one, 2013, 8(11): e80635.

43. The NCBI C++ Toolkit (https://ncbi.github.io/cxx-toolkit/) by the National Center for Biotechnology Information, U.S. National Library of Medicine; Bethesda MD, 20894 USA.

44. Henikoff S, Henikoff J G. Amino acid substitution matrices from protein blocks. Proceedings of the National Academy of Sciences, 1992, 89(22): 10915–10919.

